# Comparison of Hydroponic and Substrate-Based Methods for Screening Pea Genotypes for Resistance to *Fusarium oxysporum*

**DOI:** 10.64898/2026.10.29.762726

**Authors:** Yully Tobar, Oscar Checa, Juan Romero, Marcela Riascos, Carlos Marcillo, Eliana Revelo

**Author notes:** Corresponding author: Oscar Checa.

## Abstract

Pea (*Pisum sativum* L.) is one of the most economically and nutritionally important legumes due to its protein, vitamin, and mineral content. It also represents a source of income for producers. One of the major constraints on pea production is vascular wilt caused by the fungus *Fusarium oxysporum*. One strategy for disease management is the use of resistant varieties, which highlights the need for efficient methodologies to evaluate genotype responses to the pathogen. The objective of this study was to compare the efficiency of hydroponic and conventional substrate-based methods for evaluating the responses of pea genotypes to *F. oxysporum* under semi-controlled conditions. The hydroponic method used a modified Hoagland and Arnon nutrient solution with adjusted nutrient concentrations, whereas the conventional substrate-based method used plastic bags containing one kilogram of peat. The pathogen inoculum was obtained from pea plants affected by the disease in the municipality of Puerres, Nariño, Colombia (isolate FPUE02). In both methods, six pea genotypes and an uninoculated control were evaluated using a randomized complete block design with three replicates. Plant growth parameters, disease incidence, and disease severity were recorded. The hydroponic method showed greater sensitivity in discriminating among susceptible genotypes and facilitated the monitoring of disease progression in the roots. These results highlight the potential of the hydroponic method as a tool for the phytopathological evaluation of pea germplasm.

## Introduction

Pea (*Pisum sativum* L.) is one of the earliest crops domesticated by humans and has been an important component of the human diet since ancient times. Its origins are associated with regions such as Ethiopia, Iran, and Afghanistan, where wild pea populations can still be found. In Europe, France, Germany, the United Kingdom, and Denmark are among the main pea-producing countries. In the Americas, pea is cultivated from Canada to Chile and Argentina (Rosero and Checa 2021). In most pea-producing regions worldwide, bush-type cultivars are grown primarily for dry grain production. However, in certain regions and countries, such as Colombia, production relies mainly on climbing cultivars harvested at the green-pod stage for fresh grain consumption. This production system contributes to food security, provides an important source of income for farmers, and generates rural employment. One of the major constraints to pea production is vascular wilt caused by the fungus *Fusarium oxysporum*, which has been reported in pea-growing regions worldwide. In North Dakota, USA, Gargouri-Jbir et al. (2024) reported that Fusarium-associated root rot can cause yield losses of up to 75%, noting that *F. oxysporum* was frequently isolated from affected plants. In Egypt, El-Sharkawy et al. (2021) reported that, under favorable conditions, the aggressiveness of *F. oxysporum* can result in yield losses of up to 100%. In China, Deng et al. (2026) obtained 79 pathogenic isolates, of which 49 were identified as race 1 of the pathogen. These findings suggest that the pathogen is widely distributed and exhibits considerable variability. In Nariño, Colombia, Checa and Alpala (2016) reported production losses ranging from 30 to 100% in affected fields. These authors evaluated 18 pea lines against three isolates of *F. oxysporum* collected from the municipalities of Gualmatán, Ipiales, and Pupiales, identifying lines that exhibited moderate resistance to two or more isolates. The Andean Crops Research Group at the University of Nariño and the Colombian Agricultural Research Corporation (AGROSAVIA) maintain a collection of more than 400 pea accessions that need to be evaluated for their responses to *Fusarium oxysporum*. This evaluation aims to identify genotypes with resistance or tolerance to the pathogen that could serve as parental material in pea breeding programs in Colombia. Under controlled conditions, one of the most commonly used methods involves immersing the roots in a pathogen inoculum suspension, followed by transplanting the seedlings into bags or pots containing sterile substrate (Bani et al. 2012). However, this method requires considerable greenhouse or screenhouse space, particularly when large numbers of accessions or genotypes are evaluated. A hydroponic method may represent a promising alternative for the rapid screening of pea genotypes for resistance to *Fusarium oxysporum*, particularly when evaluating large numbers of accessions and minimizing the variability associated with soil or substrate conditions. In chickpea (Cicer arietinum), Jorben et al. (2023) developed a hydroponic system to assess the resistance of 22 genotypes to different races of *F. oxysporum* f. sp. ciceris. The genotypes were grown in Hoagland’s nutrient solution and inoculated with isolates representing five pathogen races. This method enabled the identification of differences in resistance among the genotypes evaluated. Similarly, a hydroponic protocol has been developed to assess Fusarium wilt caused by *F. oxysporum* f. sp. cubense Tropical Race 4 (TR4) in banana. This protocol employs a soil-free laboratory system using rooted plants, a conidial suspension of the pathogen, and early symptom assessment. The system was designed to be simple, portable, space- efficient, and low-maintenance, while remaining adaptable to different experimental conditions (Noordin et al., 2022). The objective of the present study was to compare the efficiency of a hydroponic method developed to evaluate the response of pea genotypes to *F. oxysporum* with that of the conventional method involving root dipping in the pathogen inoculum followed by transplantation into pots containing a substrate.

## Materials and Methods

### Study site

The study was conducted at the Botana Experimental Farm of the University of Nariño, located at an elevation of 2,811 m above sea level (m.a.s.l.), at 01°09′30″ N and 77°16′35″ W, with a mean temperature of 13 °C (AR Labs, 2025).

The effects of *Fusarium oxysporum* on pea genotypes were evaluated using two methods: a hydroponic method and a conventional substrate-based method. The experimental procedures involved the following steps.

### Isolation, purification, preservation, and propagation of *Fusarium oxysporum*

Pea plants showing symptoms of *Fusarium oxysporum* infection were collected in the municipality of Puerres, Nariño, Colombia. Sections of roots and symptomatic tissues from the first internode were aseptically plated onto potato dextrose agar (PDA) and rose bengal–chloramphenicol agar. The samples were incubated in the dark at 25 °C for eight days (Agrios, 2005). To obtain monosporic cultures, a spore suspension was prepared at a concentration of 1×1031×103 spores mL−1−1, and 2 µL aliquots were spread onto water agar. After 24 h of incubation, individual colonies were transferred to PDA and incubated at 25 °C for four days (Leslie, 2006). The monospore isolate identified as FPUE02 was preserved using the Whatman No. 1 filter paper technique. Sterile filter paper squares were placed on PDA plates and inoculated with mycelium from the monospore culture. After five days of incubation at 25 °C, the colonized filter papers were dried at 37 °C for three days and stored in cryovials at −20 °C (Fong et al., 2000).

The isolate was propagated by first reactivating it on PDA at 25 °C for four days, followed by inoculation into pea broth. The broth was prepared by boiling 300 g of fresh peas in 500 mL of distilled water for 2 h. The resulting extract was filtered and supplemented with sugar at a concentration of 30 g L⁻¹. The broth was sterilized in an autoclave at 120 °C. Once cooled, 40 mL aliquots were dispensed into sterile 8-oz containers and inoculated with mycelium from the reactivated FPUE02 isolate. The cultures were incubated at 25 °C for 30 days. Following incubation, the mycelium was homogenized by vortexing for 2 min, and the resulting suspension was filtered through two layers of sterile gauze. The inoculum concentration was determined using a Neubauer haemocytometer under a microscope at 40× magnification and adjusted with sterile distilled water to a final concentration of 5×1065×106 microconidia mL⁻¹. The isolation, purification, preservation, and propagation procedures for *Fusarium oxysporum* described above were common to both experimental methods: the hydroponic method and the conventional substrate-based method.

## Plant material preparation and inoculation

Six pea genotypes were used, comprising three commercial cultivars, ‘Sindamanoy’, ‘Obonuco San Isidro’, and ‘Alcalá’, and three breeding lines, UN7328, UN7213, and Udenar160. The latter lines have pathogen-resistant parental genotypes in their pedigrees. The six genotypes were sown in germination trays containing peat moss to obtain the seedlings required for evaluation using the two methods under comparison: the hydroponic method and the conventional substrate-based method. When the seedlings reached a height of 12 cm, they were removed from the germination trays, and their roots were washed. The root tips were then transversely cut to a length of 2 cm. The seedlings were inoculated by immersing their roots in 350 mL of a suspension containing 5×1065×106 microconidia mL⁻¹ for 30 min.

## Hydroponic method

The hydroponic system used a nutrient solution formulated based on the composition proposed by Hoagland and Arnon (1950), with modifications to nutrient concentrations. The nutrient solution contained the following components (mg L⁻¹): NO₃⁻-N, 98; P₂O₅, 52; SO₄²⁻, 64; K₂O, 282; CaO, 224; MgO, 80; and NH₄⁺-N, 0.5.

Following inoculation, the plants were placed in plastic containers filled with the nutrient solution described above. The containers were covered with aluminium foil to reduce light exposure to the roots, and an aeration system was installed to maintain oxygenation of the nutrient solution. Finally, an additional 80 mL of inoculum was added to each container.

## Conventional substrate-based method

Pea plants inoculated with *Fusarium oxysporum* were transplanted into plastic bags containing 1 kg of peat moss as the growing substrate. An additional 14 mL of inoculum was added to each bag. For the control treatment, the root tips of the seedlings were cut and immersed in pathogen-free water.

## Experimental design

A randomized complete block design (RCBD) was used for both methods (hydroponic and conventional substrate-based), with six treatments and three replicates. The treatments consisted of the six pea genotypes inoculated with *Fusarium oxysporum*, while the non-inoculated ‘Obonuco San Isidro’ cultivar was used as the control. The experimental unit consisted of 12 plants in the hydroponic method and six plants in the conventional substrate-based method.

## Variables assessed

Plant height. Plant height was measured in centimetres from the base of the plant to the tip of the uppermost leaflet. The mean plant height was then calculated for each experimental unit.

Root length. Root length was measured in centimetres from the root collar to the distal tip of the root system of each plant within the experimental unit. The mean root length was then calculated for each experimental unit. Plant height and root length were measured on seven occasions: 5, 9, 11, 16, 19, 23, and 26 days after inoculation (DAI).

Dry matter. After the final root length assessment, the plants were removed from the growing medium and placed in Kraft paper bags, with shoots and roots separated. The samples were then dried in an oven at 60 °C for 24 h, after which their dry weight was recorded in grams.

Disease incidence. In both the hydroponic and conventional substrate-based methods, plants were removed from each experimental unit, and longitudinal cuts were made in the tertiary and secondary roots, taproot, root collar, and first internode. The number of plants showing symptoms associated with *Fusarium oxysporum* was recorded for each assessed structure and divided by the total number of plants evaluated. Disease incidence was calculated using the formula described by Agrios (2005):

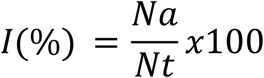

Where:

**I**: disease incidence (%).
**Na**: number of affected plants.
**Nt**: total number of plants assessed.

Disease severity. To assess disease progression based on the percentage severity, the presence of the pathogen in different root structures and the basal stem region was first evaluated. A weighting score was assigned according to the extent of pathogen progression within the plant, with lower scores assigned to pathogen presence in tertiary roots and higher scores when the pathogen was detected in the first internode. Accordingly, the weighting scores were 1 for tertiary roots, 2 for secondary roots, 3 for the taproot, 4 for the root collar, and 5 for the first internode. These scores were subsequently used to calculate disease severity (%) based on the equation proposed by García et al. (2019), adapted for the present study.

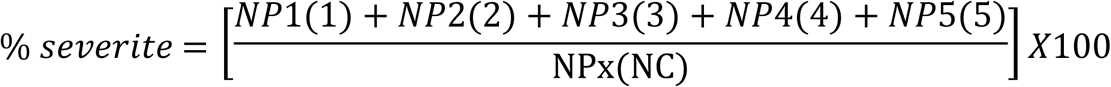

Where:

S: disease severity (%).
**NP1**: number of plants assigned a score of 1.
**NP2**: number of plants assigned a score of 2.
**NP3**: number of plants assigned a score of 3.
**NP4**: number of plants assigned a score of 4.

**NP5**: number of plants assigned a score of 5.

**NP**: total number of plants evaluated.
**NC**: number of categories in the severity scale (5)

Pathogen confirmation. The presence of the pathogen was confirmed by re-isolation from stem tissues (internodes up to the point of pathogen progression) and roots of plants exhibiting characteristic disease symptoms. This procedure helped fulfill Koch’s postulates and confirm the association between the inoculated pathogen and the symptoms observed.

## Statistical analysis

The data obtained from the evaluation of the effects of *Fusarium oxysporum* on pea genotypes using the hydroponic and conventional substrate-based methods were analysed separately by analysis of variance (ANOVA), followed by Tukey’s honestly significant difference (HSD) test for mean comparisons. Percentage data were transformed using the arcsine square-root transformation, Y= arcsin√p, where p represents the proportion expressed on a scale from 0 to 1. To address boundary values, a continuity correction was applied following the approach described by Madden et al. (2007): values of 0% were replaced with 1/(4n)1/(4n), whereas values of 100% were replaced with 1−1/(4n)1−1/(4n), where n is the number of plants evaluated per experimental unit. The arcsine square-root transformation was then applied to the corrected proportions. All statistical analyses were performed using SAS statistical software, version 9.1 (SAS Institute Inc., Cary, NC, USA).

## Results and Discussion

### Hydroponic method

The analysis of variance revealed statistically significant differences among the genotypes, including the non- inoculated control, for plant height, root length, and shoot, root, and total dry matter (Table 1). Regarding the phytopathological variables, significant differences were detected in disease incidence in the tertiary roots, secondary roots, taproot, and root collar, as well as in disease severity. In contrast, disease incidence in the first internode did not differ significantly among the treatments evaluated (Table 2).

**Table 1.**
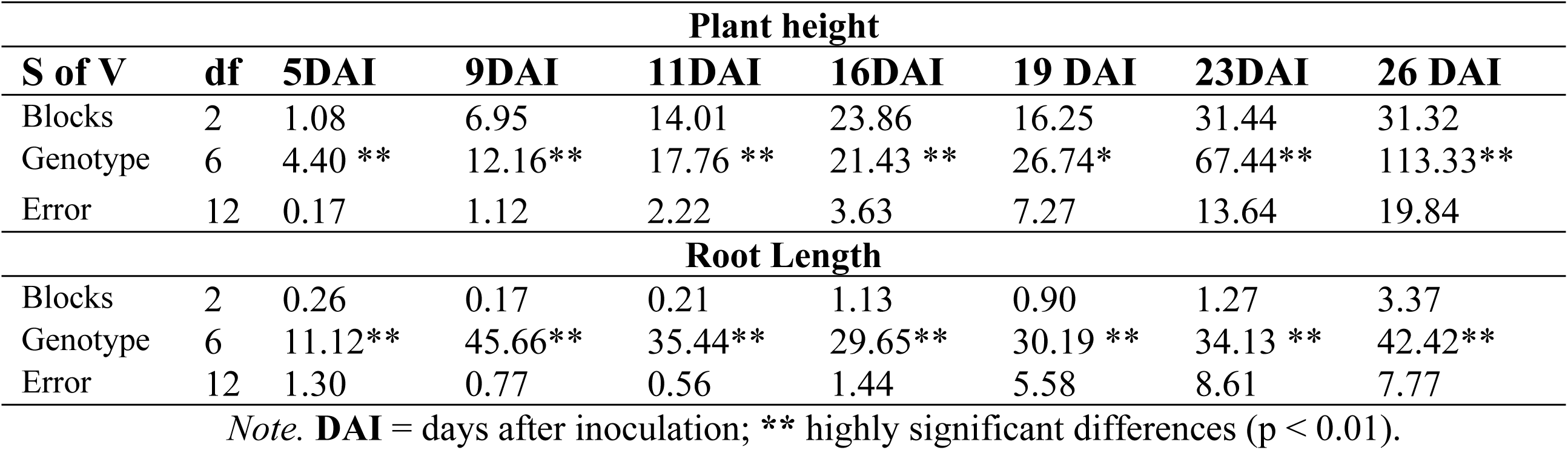
Mean squares for seven assessments of plant height and root length in six pea genotypes and a non-inoculated control following inoculation with Fusarium oxysporum using the hydroponic method.

| **Plant height** |  |  |  |  |  |  |  |  |
|---|---|---|---|---|---|---|---|---|
| **S of V** | **df** | **5DAI** | **9DAI** | **11DAI** | **16DAI** | **19 DAI** | **23DAI** | **26 DAI** |
| Blocks | 2 | 1.08 | 6.95 | 14.01 | 23.86 | 16.25 | 31.44 | 31.32 |
| Genotype | 6 | 4.40 ** | 12.16** | 17.76 ** | 21.43 ** | 26.74* | 67.44** | 113.33** |
| Error | 12 | 0.17 | 1.12 | 2.22 | 3.63 | 7.27 | 13.64 | 19.84 |
| **Root Length** |  |  |  |  |  |  |  |  |
| Blocks | 2 | 0.26 | 0.17 | 0.21 | 1.13 | 0.90 | 1.27 | 3.37 |
| Genotype | 6 | 11.12** | 45.66** | 35.44** | 29.65** | 30.19 ** | 34.13 ** | 42.42** |
| Error | 12 | 1.30 | 0.77 | 0.56 | 1.44 | 5.58 | 8.61 | 7.77 |

*Note.* **DAI** = days after inoculation; **\*\*** highly significant differences (p < 0.01).

**Table 2.** Mean squares for dry matter, disease incidence, and disease severity in six pea (Pisum sativum L.) genotypes evaluated using the hydroponic method.

| **Genotype** | **5DAI** | **9DAI** | **11DAI** | **16DAI** | **19DAI** | **23DAI** | **26DAI** |
|---|---|---|---|---|---|---|---|
| **UN7328** | 11.51 cd | 18.72 abc | 26.59 a | 36.08 a | 42.75 a | 54.97 a | 62.13 a |
| **Alcalá** | 11.40 cde | 17.45 bcd | 23.88 abc | 32.88 ab | 38.09 ab | 44.80 ab | 46.80 b |
| **Udenar160** | 13.58 a | 19.61 ab | 26.29 ab | 33.81 ab | 40.34 ab | 49.20 ab | 55.68 ab |
| **San Isidro** | 12.05 bc | 18.68 abc | 25.87 ab | 32.63 ab | 38.06 ab | 49.41 ab | 56.41 ab |
| **UN7213** | 10.25 e | 14.90 d | 21.03 c | 28.68 b | 34.45 b | 41.05 b | 45.61 b |
| **Sindamanoy** | 10.61 de | 16.04 cd | 22.05 bc | 28.76 b | 34.58 b | 42.87 b | 50.22 ab |
| **Testigo** | 13.02 ab | 20.65 a | 27.33 a | 32.76 ab | 39.23 ab | 49.06 ab | 58.08 ab |

*Note.* Different letters indicate significant differences among treatments (p ≤ 0.05).

## Plant height

At 5 days after inoculation (DAI), the genotype Udenar160 exhibited a mean plant height of 13.58 cm, exceeding those of the other genotypes evaluated, which ranged from 10.25 to 12.05 cm, except for the non- inoculated control, which reached 13.02 cm. The non-inoculated control differed significantly from UN7213, UN7328, Alcalá, and Sindamanoy, suggesting that plant growth in these genotypes may have been affected by the pathogen as early as 5 days after inoculation.

At 9 DAI, the non-inoculated control (20.65 cm) showed a mean plant height similar to those of Udenar160, Obonuco San Isidro, and UN7328, which ranged from 18.68 to 19.61 cm. In contrast, Alcalá, Sindamanoy, and UN7213 exhibited lower plant heights than the non-inoculated control, maintaining the trend observed in the first assessment at 5 DAI (Table 3). At 11 DAI, fewer differences in plant height were observed among the evaluated genotypes and the non-inoculated control. Only UN7213 and Sindamanoy, with mean heights of 21.03 and 22.05 cm, respectively, exhibited lower values than the non-inoculated control (27.33 cm) (Table 3).

**Table 3.** Comparison of mean plant heights at 5, 9, 11, 16, 19, 23, and 26 days after inoculation with Fusarium oxysporum in pea genotypes evaluated using the hydroponic method.

| **S de V** | **df** | **DMF** | **DMR** | **TDM** | **TIR** | **SIR** | **TAPIR** | **RCI** | **FII** | **SEV** |
|---|---|---|---|---|---|---|---|---|---|---|
| Blocks | 2 | 0.09 | 0.006 | 0.15 | 0.055 | 0.016 | 0.007 | 0.063 | 0.052 | 0.013 |
| Genotype | 6 | 0.17 ** | 0.010 ** | 0.24 ** | 0.268** | 0.528** | 0.647** | 0.287* | 0.056NS | 0.174** |
| Error | 12 | 0.02 | 0.001 | 0.02 | 0.016 | 0.026 | 0.007 | 0.080 | 0.020 | 0.011 |

Note. DMF = shoot dry matter; DMR = root dry matter; TDM = total dry matter; TIR = disease incidence in tertiary roots; SIR = disease incidence in secondary roots; TAPIR = disease incidence in the taproot; RCI = disease incidence at the root collar; FII = disease incidence in the first internode; SEV = disease severity; NS = not significant (p > 0.05); \* significant differences (p ≤ 0.05); \*\* highly significant diff

At 9 days after inoculation (DAI), the non-inoculated control (20.65 cm) exhibited a mean plant height comparable to those of Udenar160, Obonuco San Isidro, and UN7328, which ranged from 18.68 to 19.61 cm. In contrast, Alcalá, Sindamanoy, and UN7213 showed lower plant heights than the non-inoculated control, maintaining the trend observed in the first assessment at 5 DAI (Table 3). At 11 DAI, fewer differences in plant height were observed between the evaluated genotypes and the non-inoculated control. Only UN7213 and Sindamanoy, with mean heights of 21.03 and 22.05 cm, respectively, exhibited lower plant heights than the non-inoculated control (27.33 cm) (Table 3).

No significant differences in plant height were observed between the non-inoculated control and the evaluated genotypes at 16, 19, 23, or 26 days after inoculation (DAI). However, some differences unrelated to pathogen presence were evident. In particular, UN7328 consistently exhibited greater plant height than the UN7213 line across all seven assessments, which may be associated with the genetic background of the materials evaluated (Table 3). Inoculation with *Fusarium oxysporum* initially affected the growth of some genotypes; however, these differences diminished over time.

The lower plant heights observed at the early assessment stages (5 and 9 days after inoculation) in the genotypes Alcalá, Sindamanoy, and UN7213, compared with the non-inoculated control, suggest an early effect of the pathogen on plant growth. This finding is consistent with reports by Agrios (2005), Bani et al. (2018), Zhou et al. (2018), and Infantino et al. (2006), who indicate that *Fusarium oxysporum* colonizes the xylem, interfering with water and nutrient transport and consequently restricting plant growth during the early stages of infection. Similarly, studies in pea have shown that reductions in growth parameters, such as plant height, can occur at early stages as a consequence of pathogen-induced vascular stress (Michielse and Rep, 2009).

From 16 days after inoculation (DAI) onward, most genotypes exhibited a recovery response, possibly because the plants were able to use the nutrients available in the hydroponic solution to produce new roots, while the pathogen inoculum concentration in the solution may have decreased over time. Nevertheless, in hydroponic systems, root–pathogen interactions occur directly within the nutrient solution, potentially facilitating vascular colonization in susceptible genotypes and accelerating the onset of symptoms, such as reduced plant height and biomass. Furthermore, under hydroponic conditions, any disruption of the root system can rapidly translate into limitations in shoot development because of the plant’s strong dependence on balanced nutrient uptake (Resh, 2013).

Root length. At 5, 9, 11, and 16 days after inoculation (DAI), all evaluated genotypes exhibited lower root growth than the non-inoculated control, indicating an early reduction in root development associated with pathogen inoculation (Table 4). Among the inoculated genotypes, UN7328 and Obonuco San Isidro showed greater root lengths, whereas Alcalá and UN7213 exhibited the lowest mean values. At 19, 23, and 26 DAI, Alcalá and UN7213 continued to exhibit significantly shorter roots than the non-inoculated control, indicating that the nutrient-enriched hydroponic medium did not promote a comparable recovery response in these genotypes, which may suggest greater susceptibility to the pathogen. In contrast, at these assessment times, UN7328, Udenar160, Obonuco San Isidro, and Sindamanoy recovered root growth to levels comparable to those of the non-inoculated control (Table 4). These responses may reflect differences in the genetic backgrounds of the evaluated genotypes, potentially enabling some plants to respond to pathogen challenge by producing new roots. The availability of nutrients in the hydroponic solution may have supported this compensatory root development, although further physiological assessments would be required to confirm this explanation.

**Table 4.** Longitud de raíz en método hidropónico a los 5, 9, 11, 16, 19, 23 y 26 días después de inocular genotipos de arveja con Fusarium oxysporum.

| **Genotype** | **5DAI** | **9DAI** | **11DAI** | **16DAI** | **19DAI** | **23DAI** | **26DAI** |
|---|---|---|---|---|---|---|---|
| **UN7328** | 7.79 b | 10.40b | 15.38 b | 18.29 b | 19.84 ab | 24.81 ab | 25.50 abc |
| **Alcalá** | 9.04 b | 8.36 b | 10.97 c | 13.41 d | 14.61 b | 16.86 b | 17.76 c |
| **Udenar160** | 8.54 b | 9.91 b | 11.87 c | 14.12 cd | 17.05 ab | 21.75 ab | 24.79 abc |
| **San Isidro** | 7.33 b | 9.48 b | 14.80 b | 17.38 bc | 19.76 ab | 23.77 ab | 26.02 ab |
| **UN7213** | 7.06 b | 8.62 b | 12.05 c | 13.38 d | 14.79 b | 17.72 b | 18.20 bc |
| **Sindamanoy** | 7.52 b | 9.20 b | 12.61 c | 15.73 bcd | 19.02 ab | 23.36 ab | 24.30 abc |
| **Testigo** | 12.63 a | 19.48 a | 21.00 a | 21.98 a | 23.58a | 25.25 a | 26.87 a |

*Note.* Different letters indicate significant differences among treatments (p ≤ 0.05).

This response is consistent with expectations for hydroponic systems, in which roots are in direct contact with the nutrient solution and, consequently, with the pathogen. Under these conditions, roots become the primary site of pathogen entry and colonization. According to Laevens et al. (2024), root pathogens can affect plants more rapidly in soilless systems because of their potential to spread readily through the liquid medium. This may explain the effects on root growth observed during the first few days after inoculation.

Figure 1 shows images captured 11 days after inoculation with *Fusarium oxysporum*, illustrating the root systems of the pea genotypes evaluated under hydroponic conditions, alongside a non-inoculated control for reference. The control exhibited a healthy root system, characterized by long, light-coloured roots, indicating normal root development in the absence of the pathogen.

**Figure 1.**
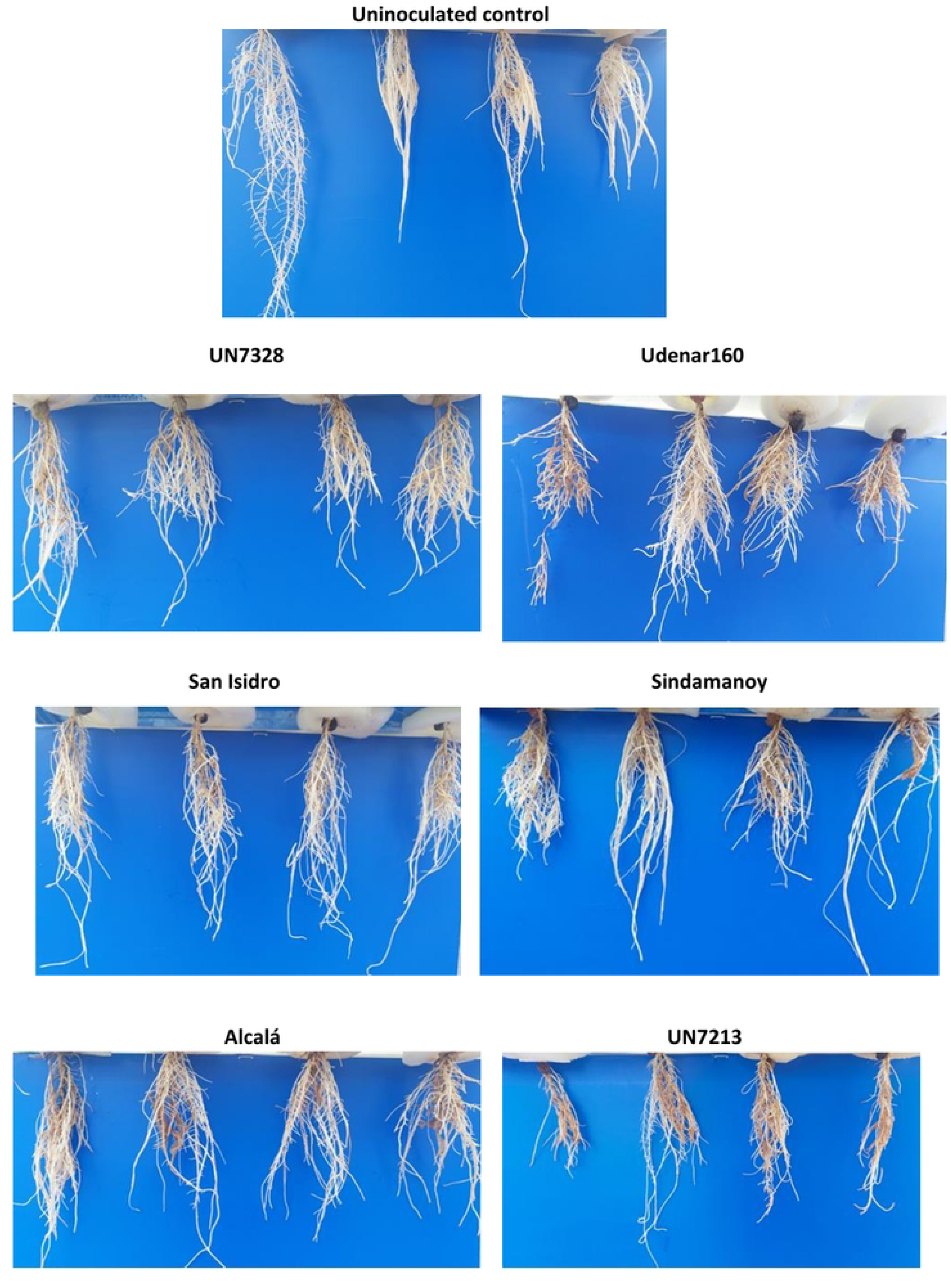
Effect of Fusarium oxysporum on root development in pea genotypes under hydroponic conditions 11 days after inoculation.

In contrast, the inoculated genotypes exhibited varying levels of root damage. Alcalá and UN7213 showed severe root deterioration, characterized by intense browning, necrosis, and a marked reduction in lateral root formation. In comparison, UN7328, Udenar160, Sindamanoy, and Obonuco San Isidro exhibited intermediate levels of damage, with part of the root structure remaining intact despite visible discoloration. However, these genotypes showed subsequent recovery in root length during later assessments. Overall, the susceptible genotypes Alcalá and UN7213 exhibited a progressive pattern of root damage, ranging from initial discoloration to advanced necrosis.

The root symptoms observed, including browning and necrosis, are characteristic of vascular wilt caused by *Fusarium oxysporum* and are consistent with previous reports in pea (Checa and Alpala, 2016; Pabón and Castaño, 2012). From a physiological perspective, browning may be interpreted as the result of the oxidation of phenolic compounds produced by the plant as part of its defense response to pathogen invasion (Agrios, 2005). However, as the infection progresses, this defense response may become insufficient, leading to necrosis of the root tissues.

Genotypes UN7328, Udenar160, San Isidro, and Sindamanoy, which exhibited lower symptom severity, may express tolerance or resistance mechanisms that favor the partial preservation of the root system in response to *Fusarium oxysporum*. According to Kraft and Pfleger (2001), genotypes with greater vegetative vigor may compensate for damage and necrosis of the primary roots through the proliferation of secondary and adventitious roots, thereby helping maintain water and nutrient uptake and reducing the risk of premature wilting.

An important aspect to highlight is the usefulness of the hydroponic method, which allowed the progression of symptoms to be observed directly and non-destructively. Unlike soil- or substrate-based systems, this method facilitates the continuous assessment of the root system under controlled conditions, enabling more precise identification of differences among genotypes and the different stages of infection. Studies such as that by Punja and Rodriguez (2018) demonstrated that, in hydroponic systems, roots infected by *F. oxysporum* exhibit a clear progression of symptoms, from early discoloration to advanced necrosis, which is consistent with the results observed in the present study.

Dry matter. The genotypes Udenar160, San Isidro, and Sindamanoy exhibited shoot and total dry matter values that were statistically similar to those of the uninoculated control. In contrast, UN7328, Alcalá, and UN7213 showed lower total dry matter accumulation, while UN7213 recorded the lowest root biomass (0.15 g). These results indicate that the effect of *F. oxysporum* on biomass accumulation varies among genotypes, with Udenar160 exhibiting the best performance (Table 5).

**Table 5.** Shoot, root, and total dry matter of six pea genotypes evaluated for their response to F. oxysporum under hydroponic conditions.

| **Genotype** | **Shoot dry matter** | **Root dry matter** | **Total dry matter** |
|---|---|---|---|
| **UN7328** | 0.93 bc | 0.16 bc | 1.09 b |
| **Alcalá** | 0.82 c | 0.24 abc | 1.07 b |
| **Udenar160** | 1.30 ab | 0.33 a | 1.63 a |
| **San Isidro** | 1.13 abc | 0.20 bc | 1.34 ab |
| **UN7213** | 0.75 c | 0.15 c | 0.90 b |
| **Sindamanoy** | 1.07 abc | 0.24 abc | 1.32 ab |
| **Testigo** | 1.40 a | 0.25 ab | 1.65 a |

*Note*. Different letters indicate significant differences among treatments (p ≤ 0.05).

When the root system is affected, the plant becomes less efficient at absorbing water and nutrients, which is rapidly reflected in reduced biomass accumulation. *Fusarium oxysporum* has been shown to invade the roots and vascular system, interfering with water and nutrient transport, thereby limiting overall plant growth and ultimately causing wilting (Perincherry et al., 2021; Zuriegat et al., 2021).

The performance of the genotypes that exhibited greater biomass accumulation (Udenar160, San Isidro, and Sindamanoy) following inoculation with *F. oxysporum* under hydroponic conditions may be associated with resistance mechanisms through which the plant restricts pathogen progression or mitigates physiological damage. Previous studies have reported that genotypes exhibiting favorable responses to F. *oxysporum* showed greater biomass accumulation, potentially associated with the activation of structural and biochemical defense mechanisms (Bani et al., 2018; Infantino et al., 2006).

Incidence. For tertiary roots, Alcalá and UN7213, with incidence rates of 57.07% and 41.67%, respectively, showed significant differences compared with the uninoculated control and the genotypes UN7328, Udenar160, San Isidro, and Sindamanoy, which exhibited incidence rates below 8.4% (Table 6).

**Table 6.**
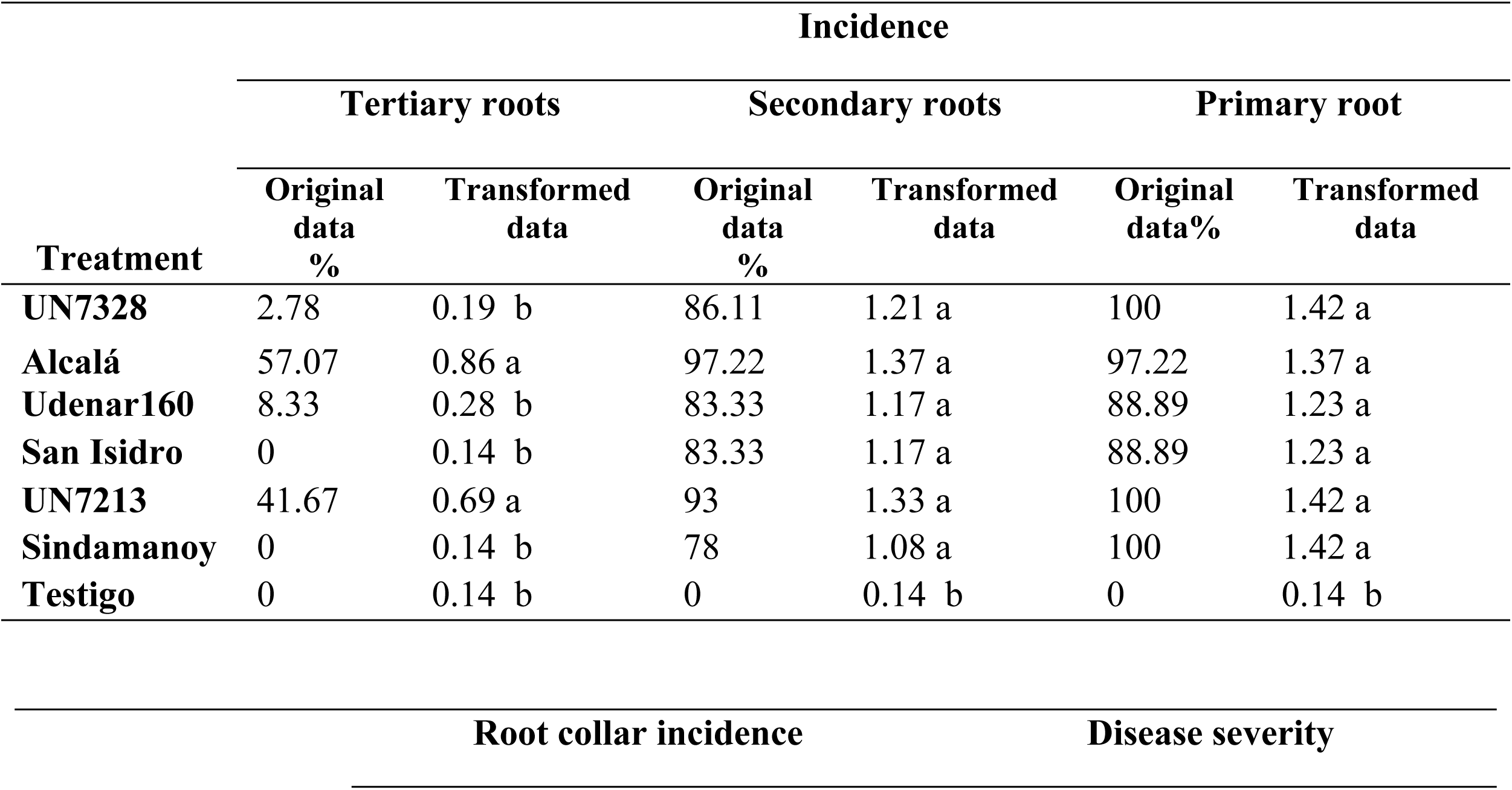
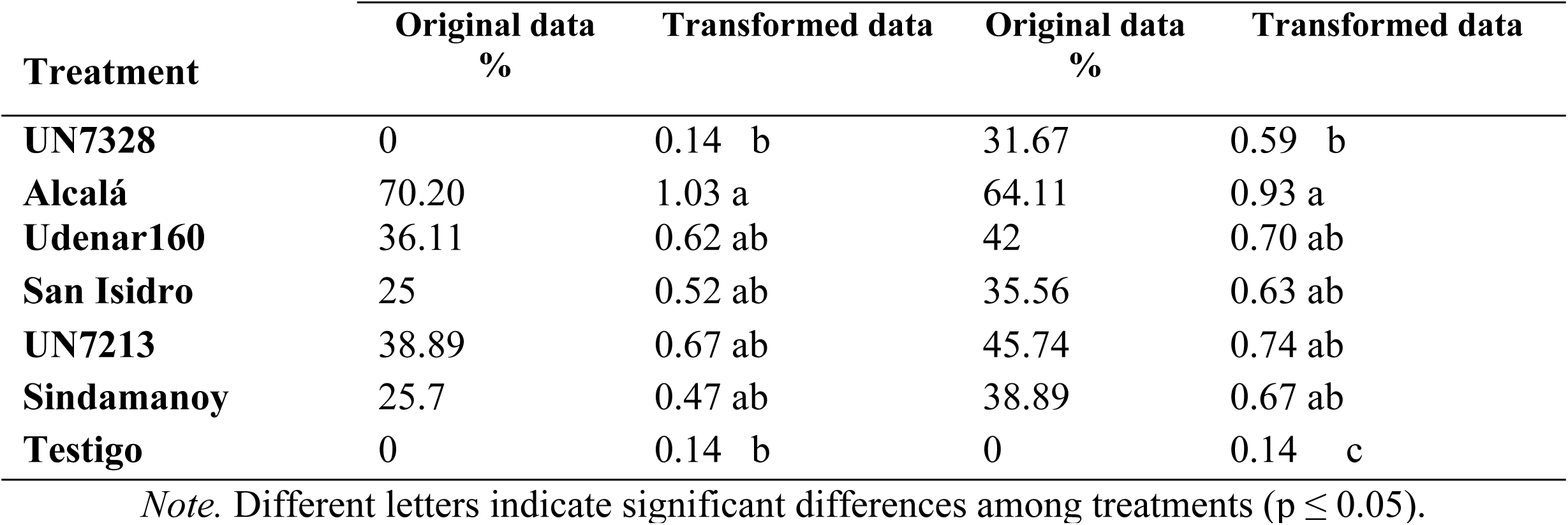
Tukey’s test for mean comparison (p ≤ 0.05) of incidence in tertiary roots, secondary roots, the primary root, and the root collar, and disease severity in pea genotypes inoculated with F. oxysporum under hydroponic conditions.

|  | Incidence |  |  |  |  |  |
|---|---|---|---|---|---|---|
|  | Tertiary roots |  | Secondary roots |  | Primary root |  |
| Treatment | Original data<br>% | Transformed data | Original data<br>% | Transformed data | Original data% | Transformed data |
| **UN7328** | 2.78 | 0.19 b | 86.11 | 1.21 a | 100 | 1.42 a |
| **Alcalá** | 57.07 | 0.86 a | 97.22 | 1.37 a | 97.22 | 1.37 a |
| **Udenar160** | 8.33 | 0.28 b | 83.33 | 1.17 a | 88.89 | 1.23 a |
| **San Isidro** | 0 | 0.14 b | 83.33 | 1.17 a | 88.89 | 1.23 a |
| **UN7213** | 41.67 | 0.69 a | 93 | 1.33 a | 100 | 1.42 a |
| **Sindamanoy** | 0 | 0.14 b | 78 | 1.08 a | 100 | 1.42 a |
| **Testigo** | 0 | 0.14 b | 0 | 0.14 b | 0 | 0.14 b |
|  | **Root collar incidence** |  | **Disease severity** |  |  |  |
| **Treatment** | **Original data<br>%** | **Transformed data** | **Original data<br>%** | **Transformed data** |  |  |
| **UN7328** | 0 | 0.14 b | 31.67 | 0.59 b |  |  |
| **Alcalá** | 70.20 | 1.03 a | 64.11 | 0.93 a |  |  |
| **Udenar160** | 36.11 | 0.62 ab | 42 | 0.70 ab |  |  |
| **San Isidro** | 25 | 0.52 ab | 35.56 | 0.63 ab |  |  |
| **UN7213** | 38.89 | 0.67 ab | 45.74 | 0.74 ab |  |  |
| **Sindamanoy** | 25.7 | 0.47 ab | 38.89 | 0.67 ab |  |  |
| **Testigo** | 0 | 0.14 b | 0 | 0.14 c |  |  |

*Note.* Different letters indicate significant differences among treatments (p ≤ 0.05).

In secondary roots and the primary root, all genotypes evaluated were affected by the pathogen, with incidence rates ranging from 78% to 100%, compared with the uninoculated control (Table 6). The genotypes UN7328, Udenar160, San Isidro, and Sindamanoy, which showed no significant pathogen incidence in tertiary roots, did not maintain this favorable response in the assessment of secondary roots and the primary root. These results suggest that the absence of detectable pathogen incidence in tertiary roots does not necessarily indicate plant resistance to the disease.

Regarding root collar incidence, only the cultivar Alcalá, with an incidence rate of 70.20%, showed significant differences compared with the uninoculated control. Furthermore, the pathogen was detected in the secondary roots and primary root of this cultivar, suggesting greater infection progression toward the upper tissues of the plant (Table 6).

Overall, the evaluated genotypes did not appear to prevent fungal entry into the roots following inoculation, which may be associated with a susceptible response. These results indicate that the hydroponic method provides suitable conditions for pathogen entry into plant roots and its progression to the first internode in more susceptible genotypes. The high incidence of infection observed may be related to the characteristics of the hydroponic system, in which the roots remain in direct contact with the nutrient solution containing the inoculum. The dispersion of microconidia in the aqueous medium promotes more uniform root exposure to the pathogen and reduces the likelihood of plants escaping infection because of heterogeneous inoculum distribution. This condition may contribute to more consistent symptom expression and facilitate the discrimination of genotype responses to *F. oxysporum* (Raizada & Nagar, 2023).

Disease severity. Under hydroponic conditions, all evaluated genotypes, with disease severity values ranging from 31.67% to 64.11%, differed significantly from the uninoculated control. Additionally, the cultivar Alcalá, which exhibited the highest disease severity (64.11%), differed significantly from UN7328 (31.67%) (Table 6). These results indicate that none of the genotypes exhibited resistance mechanisms capable of preventing pathogen progression within the roots. However, based on root length and dry matter accumulation, Udenar160, UN7328, and San Isidro exhibited the most favorable responses to pathogen attack, whereas Alcalá and UN7213 were the most susceptible genotypes.

Overall, under hydroponic conditions, the evaluated pea genotypes exhibited differential responses to inoculation with *F. oxysporum* in terms of growth, disease incidence, and severity. Although plant height tended to recover over time, root length and biomass accumulation were more severely affected, particularly in Alcalá and UN7213.

The behavior of *F. oxysporum* is consistent with that described for vascular diseases in legumes, in which resistance is quantitative; that is, plants do not prevent infection but differ in their ability to restrict pathogen development (Bani et al., 2012; Jiménez-Díaz et al., 2015). In this context, the genotype Alcalá exhibited high susceptibility and a reduced capacity to restrict vascular colonization. In contrast, UN7328 showed a more favorable response to the pathogen, possibly associated with mechanisms that limit its progression.

The greater impact on secondary roots and the primary root confirms that infection begins in the root system, as is characteristic of *F. oxysporum*, which penetrates through young roots and colonizes the xylem (Gordon, 2017; Perincherry et al., 2021; Zuriegat et al., 2021). Although pathogen progression toward the root collar and, to a lesser extent, the first internode was observed, the absence of statistically significant differences at the latter location indicates that pathogen progression into the aboveground tissues was limited. This may be associated with partial resistance mechanisms that restrict pathogen colonization (Michielse and Rep, 2009).

## Traditional Substrate-Based Method and Comparison with the Hydroponic Method

The analysis of variance (ANOVA) for the traditional substrate-based method revealed statistically significant differences among the evaluated genotypes for the following variables: plant height (in three of the seven assessment periods), root length, incidence in secondary roots, main root, root collar, and disease severity. In contrast, no statistically significant differences among genotypes were detected for shoot dry weight, root dry weight, total dry weight, or disease incidence in tertiary roots (Tables 7 and 8).

**Table 7.**
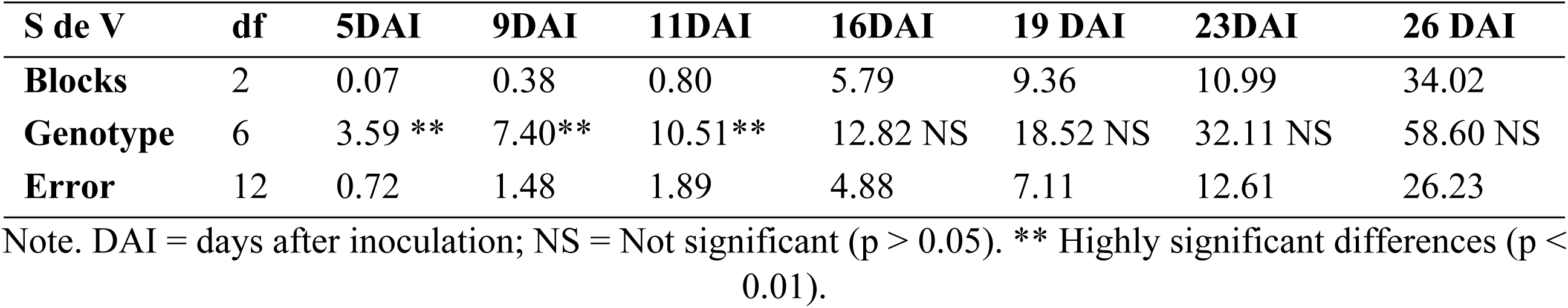
Mean squares for seven assessments of plant height in six pea genotypes following inoculation with F. oxysporum using the conventional substrate-based method.

| **S de V** | **df** | **5DAI** | **9DAI** | **11DAI** | **16DAI** | **19 DAI** | **23DAI** | **26 DAI** |
|---|---|---|---|---|---|---|---|---|
| **Blocks** | 2 | 0.07 | 0.38 | 0.80 | 5.79 | 9.36 | 10.99 | 34.02 |
| **Genotype** | 6 | 3.59 ** | 7.40** | 10.51** | 12.82 NS | 18.52 NS | 32.11 NS | 58.60 NS |
| **Error** | 12 | 0.72 | 1.48 | 1.89 | 4.88 | 7.11 | 12.61 | 26.23 |

Note. DAI = days after inoculation; NS = Not significant (p > 0.05). \*\* Highly significant differences (p < 0.01).

**Table 8.** Mean squares for growth, disease incidence, and disease severity variables in six pea genotypes evaluated for their response to F. oxysporum using the conventional substrate-based method.

| **S de V** | **df** | **RL** | **SDM** | **RDM** | **TDM** | **TRI** | **SRI** | **PRI** | **RCI** | **DS** |
|---|---|---|---|---|---|---|---|---|---|---|
| **Bloques** | 2 | 12.97 | 44.10 | 1.46 | 14.77 | 0.028 | 0.136 | 0.063 | 0.011 | 0.006 |
| **Genotipo** | 6 | 24.05** | 7.21NS | 1.43 NS | 2.72 NS | 0.130NS | 0.296* | 0.473** | 0.132* | 0.071** |
| **Error** | 12 | 5.78 | 3.23 | 0.01 | 3.17 | 0.046 | 0.069 | 0.032 | 0.028 | 0.007 |

*Note.* RL = root length; SDM = shoot dry matter; RDM = root dry matter; TDM = total dry matter; TRI = tertiary root incidence; SRI = secondary root incidence; PRI = primary root incidence; RCI = root collar incidence; DS = disease severity.
NS = Not significant. \* = Significant differences (p ≤ 0.05). \*\* = Highly significant differences (p ≤ 0.01).

## Plant Height

Under the traditional substrate-based method, at 5, 9, and 11 days post-inoculation (DPI), only the Sindamanoy genotype was affected, exhibiting a reduction in plant height compared with the non-inoculated control (Table 9). In contrast, the hydroponic method was more sensitive, revealing significant reductions in plant height in additional genotypes besides Sindamanoy. Specifically, UN7213, Alcalá, and UN7328 were affected at 5 DPI; UN7213 and Alcalá at 9 DPI; and UN7213 at 11 DPI. Over time, the differences between the two methods (hydroponic and traditional substrate-based) progressively diminished, and from 16 days post-inoculation onward, the pathogen no longer had a significant effect on plant height.

**Table 9.**
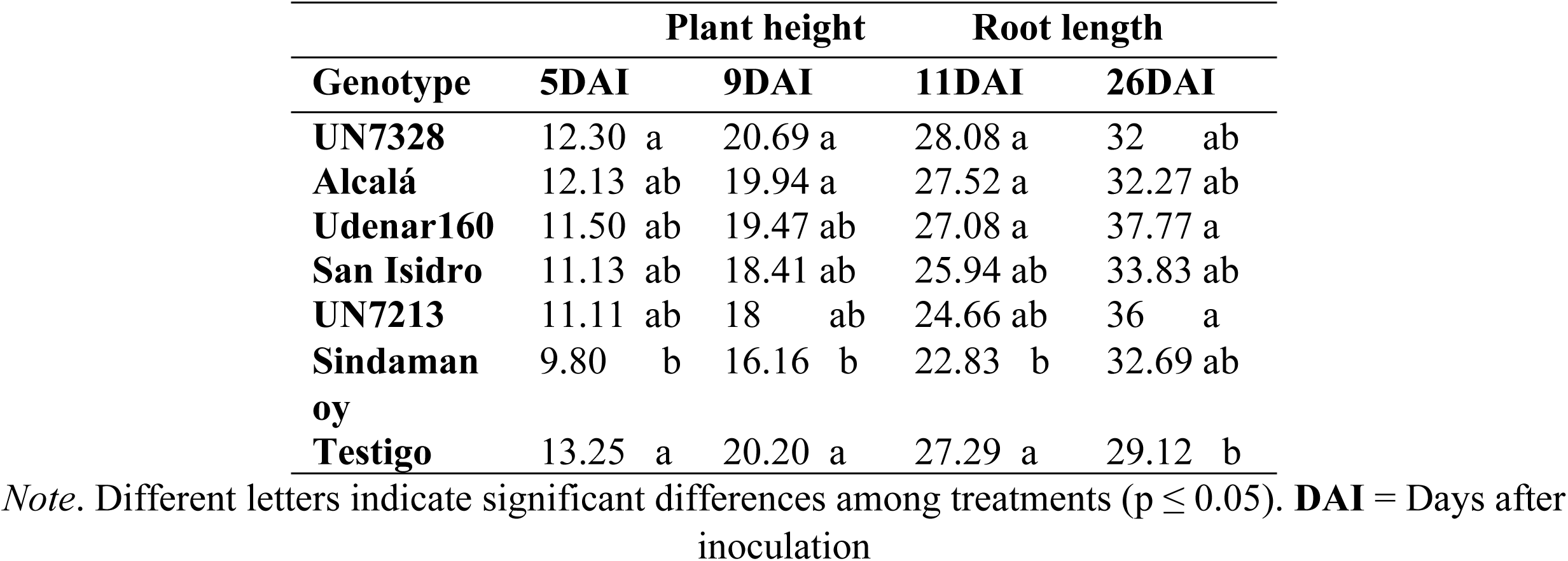
Plant height and root length under the conventional substrate-based method following inoculation of six pea genotypes with Fusarium oxysporum.

|  | **Plant height** |  | **Root length** |  |
|---|---|---|---|---|
| **Genotype** | **5DAI** | **9DAI** | **11DAI** | **26DAI** |
| **UN7328** | 12.30 a | 20.69 a | 28.08 a | 32 ab |
| **Alcalá** | 12.13 ab | 19.94 a | 27.52 a | 32.27 ab |
| **Udenar160** | 11.50 ab | 19.47 ab | 27.08 a | 37.77 a |
| **San Isidro** | 11.13 ab | 18.41 ab | 25.94 ab | 33.83 ab |
| **UN7213** | 11.11 ab | 18 ab | 24.66 ab | 36 a |
| **Sindamanoy** | 9.80 b | 16.16 b | 22.83 b | 32.69 ab |
| **Testigo** | 13.25 a | 20.20 a | 27.29 a | 29.12 b |

*Note*. Different letters indicate significant differences among treatments (p ≤ 0.05). **DAI** = Days after inoculation

*Fusarium oxysporum* is a vascular pathogen that colonizes roots and vascular bundles, disrupting water and nutrient transport. In pea, infection can reduce root growth without affecting plant height (Perincherry et al., 2021), which is consistent with the results obtained in the present study. Furthermore, the activation of biochemical defense mechanisms may mitigate damage without causing immediate morphological changes (El- Sharkawy et al., 2021). In addition, the properties of peat, including moisture retention, low compaction, porosity, and aeration, may promote physiological recovery and limit symptom expression (Hadian et al., 2025). These factors may explain why most genotypes maintained their plant height from the earliest assessments despite inoculation.

## Root length

At 26 days after inoculation (DAI), under the conventional substrate-based method, the genotypes UN7328, Alcalá, Sindamanoy, and San Isidro, with mean root lengths ranging from 32 to 33.83 cm, did not differ significantly from the uninoculated control (Table 9). These results indicate that the pathogen had no significant effect on root length in these genotypes. Udenar160 and UN7213 exceeded the uninoculated control, which is not attributable to pathogen presence but rather to genetic differences related to root length and size. In contrast, under hydroponic conditions, the genotypes Alcalá and UN7213 were affected by the pathogen, exhibiting a significant reduction in root length. These findings suggest that the conventional substrate-based method was less sensitive in discriminating among genotype responses to *F. oxysporum*. From a plant pathology perspective, the damage caused by *F. oxysporum* during the early stages of infection is primarily functional rather than structural, which may explain why root elongation can be maintained in inoculated plants of less susceptible genotypes (Michielse and Rep, 2009).

When plants interact with *F. oxysporum*, their roots may activate defense mechanisms, including cell wall lignification, which can limit pathogen colonization without necessarily impairing root growth (López-Berges et al., 2010; Pieterse et al., 2014). In pea, resistance to *F. oxysporum* has been associated primarily with the ability to restrict vascular colonization rather than with the maintenance of root growth.

## Dry matter

Under the conventional substrate-based method, shoot, root, and total dry matter did not differ significantly among genotypes (Table 8), indicating that this method did not provide sufficient sensitivity to distinguish the effects of pathogen presence in the roots on plant biomass under greenhouse conditions. In contrast, the hydroponic method identified UN7328, Alcalá, and UN7213 as the genotypes most affected in terms of root biomass.

Infection by *Fusarium oxysporum* does not necessarily result in statistically significant reductions in biomass during the early stages of plant development, as its effects on growth may vary depending on the pathogen isolate and its aggressiveness (Roots of symptom-free leguminous cover crop, 2018). Similarly, Michielse and Rep (2009) indicated that plants may continue to grow during the initial stages of pathogen colonization, possibly due to physiological compensation mechanisms. Furthermore, under controlled growing conditions and in substrates such as peat, plants may maintain biomass accumulation when adequate moisture and aeration are available, as these factors promote physiological functioning and plant growth (Taiz et al., 2015; Yafuso et al., 2019). However, Bani et al. (2018) reported significant reductions in dry matter under severe infection. These findings suggest that, in susceptible genotypes, the effects of *F. oxysporum* on biomass accumulation depend on factors such as infection intensity, plant developmental stage, and the timing of assessment.

## Incidence

Under the conventional substrate-based method, Alcalá was the only genotype that exhibited a high incidence of *F. oxysporum* in secondary roots, with a mean value of 89%, differing significantly from the uninoculated control, San Isidro (Table 10). In the primary root, all six evaluated genotypes, with incidence values ranging from 44% to 100%, differed significantly from the uninoculated control. In contrast, the hydroponic method revealed a higher incidence of the pathogen, with all six genotypes differing significantly from the uninoculated control in both secondary roots and the primary root (Table 6).

**Table 10.** Tukey’s test for mean comparison (p ≤ 0.05) of incidence in secondary roots, the primary root, and the root collar, and disease severity in pea genotypes inoculated with F. oxysporum using the conventional substrate- based method.

| **Treatment** | **Incidence** |  |  |  |  |  |  |  |
|---|---|---|---|---|---|---|---|---|
|  | **Secondary roots** |  | **Primary root** |  | **Root collar** |  | **Disease Severety** |  |
|  | % | Transformed data | % | Transformed data | % | Transformed data | % | Transformed data |
| UN7328 | 61 | 0.90 ab | 89 | 1.22 ab | 0 | 0.20 b | 28 | 0.55 a |
| Alcalá | 89 | 1.22 a | 78 | 1.08 ab | 11 | 0.34 ab | 32 | 0.60 a |
| Udenar160 | 61 | 0.90 ab | 100 | 1.36 a | 11 | 0.34 ab | 31 | 0.59 a |
| San Isidro | 42 | 0.70 ab | 67 | 0.96 ab | 0 | 0.20 b | 22 | 0.48 a |
| UN7213 | 61 | 0.90 ab | 44 | 0.72 b | 6 | 0.27 b | 21 | 0.47 a |
| Sindamanoy | 61 | 0.90 ab | 89 | 1.22 ab | 50 | 0.79 a | 40 | 0.68 a |
| Testigo | 0 | 0.20 b | 0 | 0.20 c | 0 | 0.20 b | 0 | 0.20 b |

*Note*. Different letters indicate significant differences among treatments (p ≤ 0.05).

The results obtained using the conventional substrate-based method are consistent with those reported by Checa and Alpala (2016), who evaluated *F. oxysporum* incidence in the primary root using the same method and were able to discriminate among genotypes exhibiting different levels of response to the pathogen.

Under the conventional substrate-based method, pathogen incidence at the root collar was observed only in the genotype Sindamanoy, suggesting that substrate conditions may have favored the progression of *F. oxysporum* from the primary root toward the root collar (Table 10). In contrast, under hydroponic conditions, Alcalá was the only genotype that exhibited pathogen incidence in this structure (Table 6). Together with the high incidence recorded in secondary roots and the primary root, this finding suggests greater susceptibility of this genotype to infection by the pathogen.

The incidence results across the different root structures indicate that none of the genotypes exhibited mechanisms capable of preventing pathogen entry into their roots. In pea, Checa and Alpala (2016) reported an incidence of *F. oxysporum* of 70%. In controlled pathogenicity assays conducted in inoculated substrate without biological control agents. The proportion of plants showing no evidence of pathogen presence may be attributable to random escapes during inoculation or to a small number of plants capable of preventing fungal penetration into their roots.

## Disease severity

Under the traditional substrate-based method, disease severity results, which reflect disease progression through the different structures of the root system, showed that all six genotypes inoculated with *F. oxysporum* differed significantly from the uninoculated control. These findings suggest that the *F. oxysporum* isolate FPUE02 used in this study was pathogenic and affected the roots of the genotypes evaluated. However, disease progression ranged from low to moderate, with severity values between 21 and 40% (Table 11). Similar results were obtained using the hydroponic method, although severity values were higher, with mean values ranging from 31.67 to 64.11% (Table 6).

Šišić et al. (2018) used a root rot severity scale in which symptom progression was assessed primarily on the basis of discoloration and deterioration of the taproot, ranging from the absence of symptoms to the complete loss of this structure. The authors also reported substantial variability in the aggressiveness of *Fusarium* isolates evaluated in pea, with severity indices ranging from 15 to 50, demonstrating the usefulness of root assessments for differentiating plant responses to the pathogen. In the present study, severity values were lower than those reported by these authors, particularly under the traditional substrate-based method. This difference may be attributable to variations in the assessment methods used, as well as potential differences in the genetic diversity of the genotypes evaluated.

Overall, the hydroponic method proved more discriminating, enabling the identification of a greater number of genotypes with susceptible reactions based on plant height, dry matter content, and root disease severity than the traditional substrate-based method. This is advantageous when evaluating large numbers of genotypes within shorter time frames and in limited space. Furthermore, the hydroponic method allows continuous observation of disease progression and facilitates pathogen inactivation, thereby improving experimental biosafety. Taken together, these characteristics make the hydroponic method an efficient tool for the rapid, large-scale screening of germplasm in plant breeding programs.

## Funding

This work was supported by MINCIENCIAS Colombia and the Office of Research, Graduate Studies, and International Relations of the Universidad de Nariño.

## Acknowledgements

The authors thank the members of the Andean Crops Research Group (GRICAND), Faculty of Agricultural Sciences, Universidad de Nariño, and the Colombian Agricultural Research Corporation (AGROSAVIA) for their technical support and collaboration in establishing and evaluating the greenhouse trials. The authors also acknowledge the Ministry of Science, Technology and Innovation of Colombia (Minciencias) for providing financial support for this research.

## Author Contributions

Yully Tobar: Investigation, Methodology and Writing – original draft.

Oscar Checa: Project administration, Software, Formal analysis and Writing – review & editing.

Juan Vicente Romero: Software, Conceptualization, Supervision.

Marcela Riascos: Conceptualization, Writing – review & editing and Supervision. Carlos Marcillo: Data curation and Methodology.

Eliana Revelo: Investigation and Methodology.

## Conflict of Interest

The authors declare no conflicts of interest related to the content of this article.

## Notes

### Competing Interest Statement

The authors have declared no competing interest.

## References

1. Agrios GN (2005) Plant pathology, 5th edn. Elsevier Academic Press, Burlington, MA. 10.1016/C2009-0-02037-6.

2. AR Labs. (2025). Altímetro preciso (Versión 1.0) [Aplicación móvil]. Google Play Store. google.com.

3. Bani M, Cimmino A, Evidente A, Rubiales D, Rispail N (2018) Pisatin involvement in the variation of inhibition of *Fusarium oxysporum* f. sp. pisi spore germination by root exudates of Pisum spp. germplasm. Plant Pathology 67:1046–1054. 10.1111/ppa.12813.

4. Bani, M., Rubiales, D., & Rispail, N. (2012). A detailed evaluation method to identify sources of quantitative resistance to *Fusarium oxysporum* f. sp. pisi race 2 within a *Pisum* spp. germplasm collection. Plant Pathology, 61(3), 532–542. 10.1111/j.1365-3059.2011.02537.x.

5. Checa, O., & Alpala, F. (2016). Reacción de 18 líneas de arveja (*Pisum sativum* L.) a tres cepas de *Fusarium oxysporum* f. sp. *pisi* en Nariño, Colombia. Temas Agrarios, 21(1), 28–43. 10.21897/rta.v21i1.868.

6. Deng, D., Wu, W., Duan, C., Xue, R., Sun, S., & Zhu, Z. (2026). Characterization of *Fusarium oxysporum* species complex isolates associated with pea Fusarium wilt in China. Plant Disease. 10.1094/PDIS-12-25-2438-RE.

7. El-Sharkawy, H. H. A., Abbas, M. S., Soliman, A. S., Ibrahim, S. A., & El-Nady, I. A. I. (2021). Synergistic effect of growth-promoting microorganisms on bio-control of *Fusarium oxysporum* f. sp. *pisi*, growth, yield, physiological and anatomical characteristics of pea plants. Pesticide Biochemistry and Physiology, 178, 104939. 10.1016/j.pestbp.2021.104939.

8. Fong, Y. K., Anuar, S., Lim, H. P., Tham, F. Y., & Sanderson, F. R. (2000). A modified filter paper technique for long-term preservation of some fungal cultures. Mycologist, 14(3), 127–130. 10.1016/S0269-915X(00)80090-7.

9. García, S., Pérez, O., Marín, V., & Cibrián, D. (2019). Incidencia y severidad por Fusarium spp., y umbral de daño ocasionado por moscas fungosas Bradysia y Lycoriella [Material de consulta]. Sistema Nacional de Vigilancia Epidemiológica Fitosanitaria. cnf.gob.mx.

10. Gargouri Jbir, T., Zitnick-Anderson, K., Pasche, J. S., & Kalil, A. K. (2024). Characterization of *Fusarium oxysporum* f. sp. *pisi* Associated with Root Rot of Field Pea in North Dakota and the Effects of Temperature on Aggressiveness. Plant Disease, 108(2), 365–374. DOI: 10.1094/PDIS-05-23-0908-RE.

11. Gordon TR (2017) *Fusarium oxysporum* and the *Fusarium* Wilt Syndrome. Annual Review of Phytopathology 55:23–39. 10.1146/annurev-phyto-080615-095919.

12. Hadian S, Smith DL, Suproniene S (2025) Genomic and phenotypic profiling of an Artemisia endophyte: dual biostimulant and biocontrol activities in pea under *Fusarium* stress. Frontiers in Microbiology 16:1643204. 10.3389/fmicb.2025.1643204.

13. Infantino A, Kharrat M, Riccioni L, Coyne CJ, McPhee KE, Grünwald NJ (2006) Screening techniques and sources of resistance to root diseases in cool season food legumes. Euphytica 147:201–221. 10.1007/s10681-006-6963-z.

14. Jiménez-Díaz RM, Castillo P, Jiménez-Gasco MM, Landa BB, Navas-Cortés JA (2015) *Fusarium* wilt of chickpeas: Biology, ecology and management. Crop Protection 73:16–27. 10.1016/j.cropro.2015.02.023.

15. Jorben, J., Rao, A., Nagappa Chowluru, S., Tomar, S., Kumar, N., Bharadwaj, C., Siddanagowda Patil, B., & Soren, K. R. (2023). Identification of multi-race *Fusarium* wilt resistance in chickpea (Cicer arietinum L.) using rapid hydroponic phenotyping. Phytopathologia Mediterranea, 62(1), 3–15. 10.36253/phyto-13352.

16. Jorben, J. et al. (2023). Identification of multi-race *Fusarium* wilt resistance in chickpea (Cicer arietinum L.) using rapid hydroponic phenotyping. Phytopathologia Mediterranea, 62(1), 3–15. DOI: 10.36253/phyto-13352.

17. Kraft JM, Boge W (2001) Root characteristics in pea in relation to compaction and *Fusarium* root rot. Plant Disease 85(9):936–940. 10.1094/PDIS.2001.85.9.936.

18. Laevens GCS, Dolson WC, Drapeau MM, Telhig S, Ruffell SE, Rose DM, Glick BR, Stegelmeier AA (2024) The good, the bad, and the fungus: insights into the relationship between plants, fungi, and oomycetes in hydroponics. Biology 13(12):1014. 10.3390/biology13121014.

19. Leslie JF, Summerell BA (2006) The Fusarium laboratory manual. Blackwell Publishing, Ames, IA, USA. 10.1002/9780470278376.

20. López-Berges, M. S., Rispail, N., Prados-Rosales, R. C., & Di Pietro, A. (2010). A nitrogen response pathway regulates virulence functions in *Fusarium oxysporum* via the protein kinase TOR and the bZIP protein MeaB. The Plant Cell, 22(7), 2459–2475. 10.1105/tpc.110.075937.

21. Michielse CB, Rep M (2009) Pathogen profile update: *Fusarium oxysporum*. Molecular Plant Pathology 10(3):311–324. 10.1111/j.1364-3703.2009.00538.x.

22. Noordin, N., Abu Hassan, A., Mohd Faisol Mahadevan, A. N. M., Ahmad, Z., & Ariffin, S. (2022). Lab-Based Screening Using Hydroponic System for the Rapid Detection of *Fusarium* Wilt TR4 Tolerance/Resistance of Banana. En J. Jankowicz-Cieslak & I. L. Ingelbrecht (Eds.), Efficient Screening Techniques to Identify Mutants with TR4 Resistance in Banana (pp. 79–95). Springer, Cham. 10.1007/978-3-662-64915-2_6.

23. Pabón-Villalobos J, Castaño-Zapata J (2012) Manejo de la pudrición radical de la arveja (*Pisum sativum* L.) causada por *Fusarium oxysporum* Schlechtend.:Fr. Agronomía 20(2):37–44.

24. Punja, Z. K., & Rodriguez, G. (2018). Fusarium and Pythium species infecting roots of hydroponically grown marijuana (Cannabis sativa L.) plants. Canadian Journal of Plant Pathology, 40(4), 498–513. 10.1080/07060661.2018.1535466.

25. Roots of symptom-free leguminous cover crop and living mulch species harbor diverse *Fusarium* communities that show highly variable aggressiveness on pea (*Pisum sativum*). (2018). PLOS ONE. 10.1371/journal.pone.0191969.

26. Raizada M, Nimmy MS, Nagar R (2023) A scalable hydroponic-based method for screening isolates of chickpea *Fusarium* wilt pathogen, Fusarium oxysporum f. sp. ciceris. protocols.io. 10.17504/protocols.io.5qpvormyxv4o/v1.

27. Resh HM (2013) Hydroponic food production: a definitive guidebook for the advanced home gardener and the commercial hydroponic grower, 7th edn. CRC Press, Boca Raton, FL. 10.1201/b12500.

28. Pieterse CMJ, Zamioudis C, Berendsen RL, Weller DM, Van Wees SCM, Bakker PAHM (2014) Induced Systemic Resistance by Beneficial Microbes. Annual Review of Phytopathology 52:347–375. 10.1146/annurev-phyto-082712-102340.

29. Perincherry L, Urbaniak M, Pawłowicz I, Kotowska K, Waśkiewicz A, Stępień Ł (2021) Dynamics of *Fusarium* mycotoxins and lytic enzymes during pea plants’ infection. International Journal of Molecular Sciences 22(18):9888. 10.3390/ijms22189888.

30. Rosero-Lombana V, Checa-Coral O (2021) Morphological characterization and hierarchical classification of 40 bush pea genotypes (*Pisum sativum* L.). Revista Colombiana de Ciencias Hortícolas 15(2):e12078. 10.17584/rcch.2021v15i2.12078.

31. Šišić, A., Baćanović-Šišić, J., Karlovsky, P., Wittwer, R., Walder, F., Campiglia, E., Radicetti, E., Friberg, H., Baresel, J. P., & Finckh, M. R. (2018). Roots of symptom-free leguminous cover crop and living mulch species harbor diverse *Fusarium* communities that show highly variable aggressiveness on pea (*Pisum sativum*). PLOS ONE, 13(2), Artículo e0191969. 10.1371/journal.pone.0191969.

32. Taiz L, Zeiger E, Møller IM, Murphy A (2015) Plant Physiology and Development. 6th ed. Sinauer Associates, Sunderland, MA, USA. ISBN: 978-1-60535-255-8.

33. Yafuso EJ, Fisher PR, Bohórquez AC, Altland JE (2019) Water and air relations in propagation substrates. HortScience 54(11):2024–2030. 10.21273/HORTSCI14145-19.

34. Zhou, S., Bae, J. S., Bergstrom, G. C., & Jander, G. (2018). *Fusarium* graminearum-induced shoot elongation and root reduction in maize seedlings correlate with later seedling blight severity. Plant Direct, 2(7), e00075. 10.1002/pld3.75.

35. Zuriegat Q, Zheng Y, Liu H, Wang Z, Yun Y (2021) Current progress on pathogenicity-related transcription factors in *Fusarium oxysporum*. Molecular Plant Pathology 22(7):882–895. 10.1111/mpp.13068.

